# Disruption of a BTB-ZF transcription factor causes female sterility and melanization in the larval body of the silkworm, *Bombyx mori*

**DOI:** 10.1101/2023.04.01.535244

**Authors:** Kenta Tomihara, Takashi Kiuchi

## Abstract

The *dilute black* (*bd*) of the silkworm *Bombyx mori* is a recessive mutant that produces a grayish-black color in the larval integument, instead of the characteristic white color found in wild-type larvae. In addition, eggs produced by *bd* females are sterile due to a deficiency in the micropylar apparatus. We identified candidate genes responsible for the *bd* phenotype using publicly available RNA-seq data. One of these candidate genes was homologous to the *maternal gene required for meiosis* (*mamo*) of *Drosophila melanogaster*, which encodes a broad-complex, tramtrack, and bric-à-brac-zinc finger (BTB-ZF) transcription factor essential for female fertility. In three independent *bd* strains, the expression of the *B. mori mamo* (*Bmmamo*) was downregulated in the larval integument. Using a CRISPR/Cas9-mediated knockout strategy, we found that *Bmmamo* knockout mutants exhibit a grayish-black color in the larval integument and female infertility. Moreover, larvae obtained from the complementation cross between *bd/+* mutants and heterozygous knockouts for the *Bmmamo* also exhibited a grayish-black color, indicating that *Bmmamo* is responsible for the *bd* phenotype. Gene expression analysis using *Bmmamo* knockout mutants suggested that the BmMamo protein suppresses the expression of melanin synthesis genes. Previous comparative genome analysis revealed that the *Bmmamo* was selected during silkworm domestication, and we found that *Bmmamo* expression in the larval integument is higher in *B. mori* than in the wild silkworm *B. mandarina*, suggesting that the *Bmmamo* is involved in domestication-associated pigmentation changes of the silkworm.

**Graphical Abstract:** 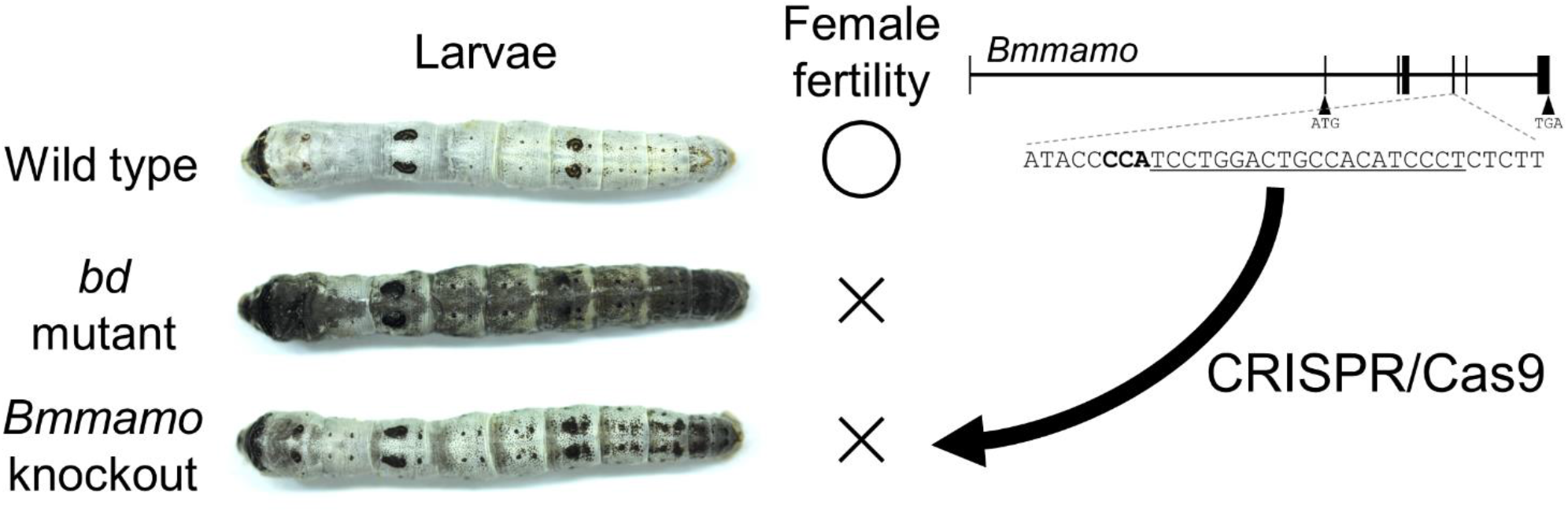

## 1. Introduction

The silkworm, *Bombyx mori*, was domesticated over 5000 years ago from its wild ancestor, *B. mandarina*. Wild-type *B. mori* larvae commonly have a white color, while *B. mandarina* larvae have a black melanin pigmentation pattern on the dorsal side (Figure 1A-C). The white color of *B. mori* is suitable in sericulture because farmers can easily identify larvae in mulberry leaves and twigs (Figure 1A). Conversely, in *B. mandarina*, the pigmentation pattern is important for mimicking bird droppings (Figure 1B) and mulberry twigs (Figure 1C). Additionally, in *B. mandarina*, melanin pigmentation is also important in terms of preventing ultraviolet damage (Hu et al., 2013).

**Figure 1.**
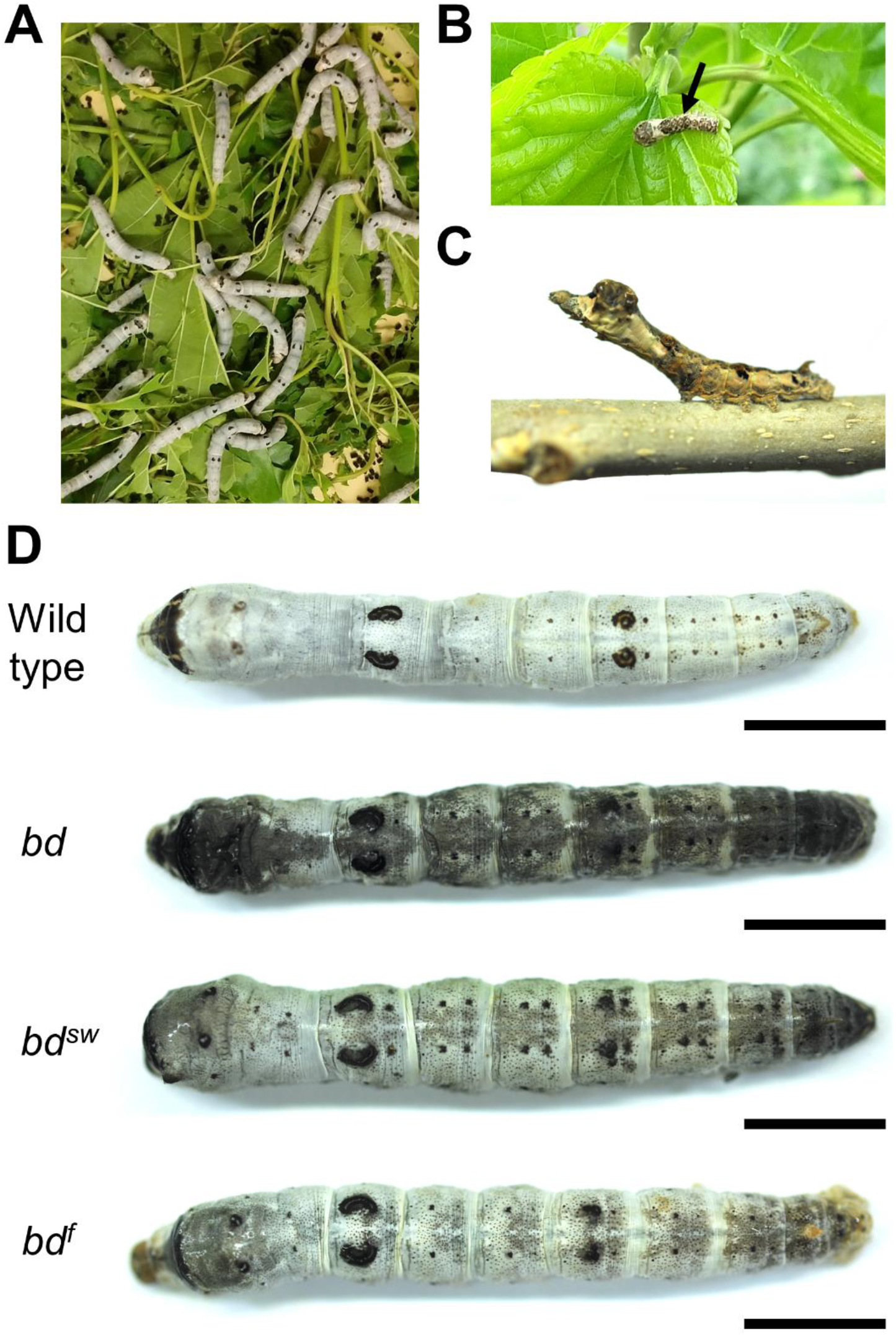
Larval phenotypes of wild-type *B. mori* and *B. mandarina*, and *bd* mutant larvae. (A) Fifth instar larvae of *B. mori*. Wild-type white *B. mori* larvae can be easily distinguished from mulberry twigs and leaves. (B) A young instar larva of *B. mandarina* mimicking bird dropping. (C) A fourth instar larva of *B. mandarina* mimicking a mulberry twig. (D) Larval phenotypes of wild-type (p50T), *bd*, *bd^sw^*, and *bd^f^* larvae in the fifth instar (day 3). Scale bars: 1cm.

In the past decade, *B. mori* mutants have been used to identify candidate genes involved in color variation as a result of selection during domestication. Yoda et al. (2014) used positional cloning to identify the gene responsible for diverse coloration pattern in larvae of *B. mori* strains (*apontic like*; *apt-like*), which was afterward found to be involved in the pigmentation pattern variation between *B. mori* and *B. mandarina* larvae (Tomihara et al., 2022). Wang et al. (2022) also performed positional cloning and found that the abnormal expression of the transcription factor gene *SoxD* is responsible for *B. mori* larval color mutation *Ursa* (*U*). *SoxD* was likely under positive selection during silkworm domestication (Wei et al., 2011), suggesting that it is also involved in color differences between larvae of *B. mori* and *B. mandarina*. A positional cloning strategy identified the *Aspartate decarboxylase* (*ADC*) gene responsible for the *black pupa* (*bp*) mutant phenotype in *B. mori* (Dai et al., 2015). The black color in *B. mandarina* pupae is likely controlled by the *bp locus* (Harizuka, 1947; Hashiguchi, 1964), and the *ADC* gene was a potential target of selection during silkworm domestication (Yu et al., 2011).

The *dilute black* (*bd*) locus has at least three alleles, *bd* (Sasaki, 1941), *chimney sweep* (*bd^sw^*) (Chikushi et al., 1972), and *dilute black fertile* (*bd^f^*) (Shimizu and Matsuno, 1978). *bd*, *bd^sw^*, and *bd^f^* larvae exhibit grayish-black coloration, while wild-type larvae have white color (Figure 1C). Moreover, *bd* and *bd^sw^* females are sterile, while *bd^f^* females are fertile. The micropylar apparatus, which plays an essential role during fertilization, is defective in oviposited eggs of *bd* or *bd^sw^*females, whereas it is normal in eggs of *bd^f^* females (Kawaguchi et al., 2006; Sakanoshita, 1970). This deficiency in the micropylar apparatus leads to female sterility in *bd* and *bd^sw^* mutants. Wu et al. (2016) performed a targeted transcriptomic analysis using integument tissue samples of *bd* mutants; however, genes associated with the mutant phenotype were not identified.

In the present study, a gene encoding a broad-complex, tramtrack, and bric-à-brac-zinc finger (BTB-ZF) transcription factor was identified to be responsible for the *bd* mutant phenotype. This gene regulates the transcription of melanin synthesis genes and may have experienced selection during silkworm domestication.

## 2. Materials and Methods

### 2.1 Insects

In this study, p50T and Sakado, wild-type strains of *B. mori* and *B. mandarina*, respectively, were maintained in our laboratory. *bd* mutant strains, namely, k40 (*bd*), k42 (*bd^sw^*), and k45 (*bd^f^*), were obtained from the Kyushu University with support of the National Bioresource Project (http://silkworm.nbrp.jp/). Larvae were reared on fresh mulberry leaves or with an artificial diet (Silkmate PS [NOSAN]) under a 12 h light and 12 h dark cycle at 25°C with exceptions as described below.

### 2.2 RNA-sequence (RNA-seq) data analysis

Previously published RNA-seq data (Wu et al., 2016) were used. Low-quality reads and adapter sequences were removed from raw reads using Trim Galore! (https://www.bioinformatics.babraham.ac.uk/projects/trim_galore/). Trimmed reads were mapped to the *B. mori* reference genome (Kawamoto et al., 2019) by the STAR 2.7.10 (Dobin et al., 2013), and then BAM files were converted to VCF using GATK 4.3.0 (McKenna et al., 2010). The code used in mapping and variant calling can be obtained from https://github.com/ketomihara/bd_codes. The number of heterozygous single nucleotide polymorphisms (SNPs) was calculated using a custom R script (https://github.com/ketomihara/bd_codes). Count data and transcripts per million values for each gene were calculated using RSEM (Li and Dewey, 2011). The differential expression analysis was performed using the edgeR R package (Robinson et al., 2010).

### 2.3 RNA extraction, RT-PCR and qPCR

Total RNA was extracted from the larval integument using TRIzol Reagent (Invitrogen). Reverse transcription was performed using the oligo (dT) primer and AMV reverse transcriptase contained in the TaKaRa RNA PCR kit (TaKaRa). The RT-PCR was performed using KOD One PCR Master Mix (TOYOBO) under the following conditions: initial denaturation at 94°C for 1 min, and then 35 cycles at 98°C for 10 s, 60°C for 5 s, and 68°C for 5 s/kb. PCR products were sequenced by the FASMAC sequencing service. The qPCR analysis was performed using the KAPA SYBR FAST qPCR Master Mix Kit (Kapa Biosystems) and amplifications were carried out in the Step One Plus Real-Time PCR System (Applied biosystems). Sequences of forward and reverse primers are listed in Table S1.

### 2.4 Phylogenetic analysis

Protein sequences of *B. mori*, *Trichoplusia ni*, *Drosophila melanogaster*, *Musca domestica, Aedes aegypti, Apis mellifera*, *Bombus terrestris*, *Tribolium castaneum*, *Nicrophorus vespilloides*, *Acyrthosiphon pisum*, *Cimex lectularius*, and *Zootermopsis nevadensis* were collected from the NCBI genome database (https://www.ncbi.nlm.nih.gov/genome/). For each protein dataset, proteins with both the Pfam domains “BTB/POZ domain” (PF00651) and “Zinc finger, C2H2 type” (PF00096) were searched using HMMsearch (Eddy, 2009) with an E-value cutoff of 1e-10. The redundant sequences were then removed by ENBOSS skipredundant (http://emboss.open-bio.org/rel/rel6/apps/skipredundant.html) at a sequence identity threshold cutoff of 60%. Note that proteins encoded by recently duplicated genes may be removed in this process. The proteins were then filtered against the BmMamo protein using the BLASTP program with an E-value cutoff of 1e-10. All the sequences were aligned using the MUSCLE program (Edgar, 2004), and a phylogenetic tree was constructed by MEGA X software (Kumar et al., 2018) using the neighbor joining method based on Jones–Taylor–Thornton (JTT) matrix model. Positions with less than 80% site coverage were eliminated in the construction of the phylogenetic tree. The topology was selected with superior log likelihood value with 1000 bootstrap replicates.

### 2.5 CRISPR/Cas9-mediated knockout

p50T larvae were reared under a 20 h light and 4 h dark daily cycle at 25°C to produce non-diapausing eggs (Kogure, 1933). We designed a unique CRISPR RNA (crRNA) target site for the *B. mori* genome using CRISPRdirect (Naito et al., 2015) (Table S1). A mixture consisting of crRNA (200 ng/µL; FASMAC), trans-activating crRNA (tracrRNA) (200 ng/µL; FASMAC), and Cas9 Nuclease protein NLS (600 ng/µL; NIPPON GENE) in injection buffer (100 mM KOAc, 2 mM Mg(OAc)_2_, 30 mM HEPES-KOH; pH 7.4) was injected into non-diapausing eggs within a period from 2 h to 6 h after oviposition (Kiuchi and Katsuma, 2022; Yamaguchi et al., 2011). Finally, embryos were incubated at 25°C in a humidified Petri dish until hatching.

Generation 0 (G_0_) moths were crossed with wild-type moths to obtain generation 1 (G_1_) individuals. Then the genomic DNA was extracted from adult G_1_ moth legs using the HotSHOT method (Truett et al., 2000). The PCR was performed using the KOD One PCR Master Mix and primers listed in Table S1, under the conditions as described above. Mutations at the targeted site were detected by the heteroduplex mobility assay using the MultiNA microchip electrophoresis system (SHIMADZU) with the DNA-500 reagent kit (Ansai et al., 2014; Ota et al., 2013). G_1_ individuals with a heterozygous knockout mutation were intercrossed with each other to generate homozygous knockout generation 2 (G_2_) individuals. PCR products of G_2_ adults were sequenced to confirm the generation of mutants carrying three mutations, namely, *Bmmamo^Δ4^*, *Bmmamo^Δ15A^*, and *Bmmamo^Δ15B^*. *Bmmamo^Δ4^* knockout mutants were maintained for further experiments as they carried a frameshift and premature stop mutation. *Bmmamo^Δ4^* females are infertile and therefore we had to maintain this strain by crossing heterozygous *Bmmamo^Δ4/+^* females to homozygous *Bmmamo^Δ4^*males.

## 3. Results

### 3.1 *In silico* identification of candidate genes from RNA-seq data

The *bd* mutation was first found in Gunze-Ao strain (Sasaki, 1941). Since *bd/bd* females are sterile, the *bd* strain has been maintained by crossing *bd/+* females with *bd/bd* males. Thus, in the *bd* strain, *+^bd^*-linked chromosomes have been transmitted through heterozygous females without recombination over generations because crossing over is absent in oocytes of *B. mori* (Tanaka, 1913). If the *bd* strain has been maintained by sib-mating since its establishment, the genetic backgrounds of the *+^bd^*-linked and *bd*-linked chromosomes are expected to be heterozygous due to an accumulation of independent mutations, while the other chromosomes are highly homozygous (Figure 2A). Alternatively, if the *bd* strain has been crossed with a strain other than Gunze-Ao and subsequently inbred, the origins of the *bd*-linked and *+^bd^*-linked chromosomes may be different (Figure 2B). In this case, it is possible that the high density of heterozygous SNPs between *+^bd^*-linked and *bd*-linked chromosomes is limited to the region closely linked to the *bd* mutation, while heterozygosity of the other chromosomes depends on the degree of inbreeding (Figure 2B).

**Figure 2.**
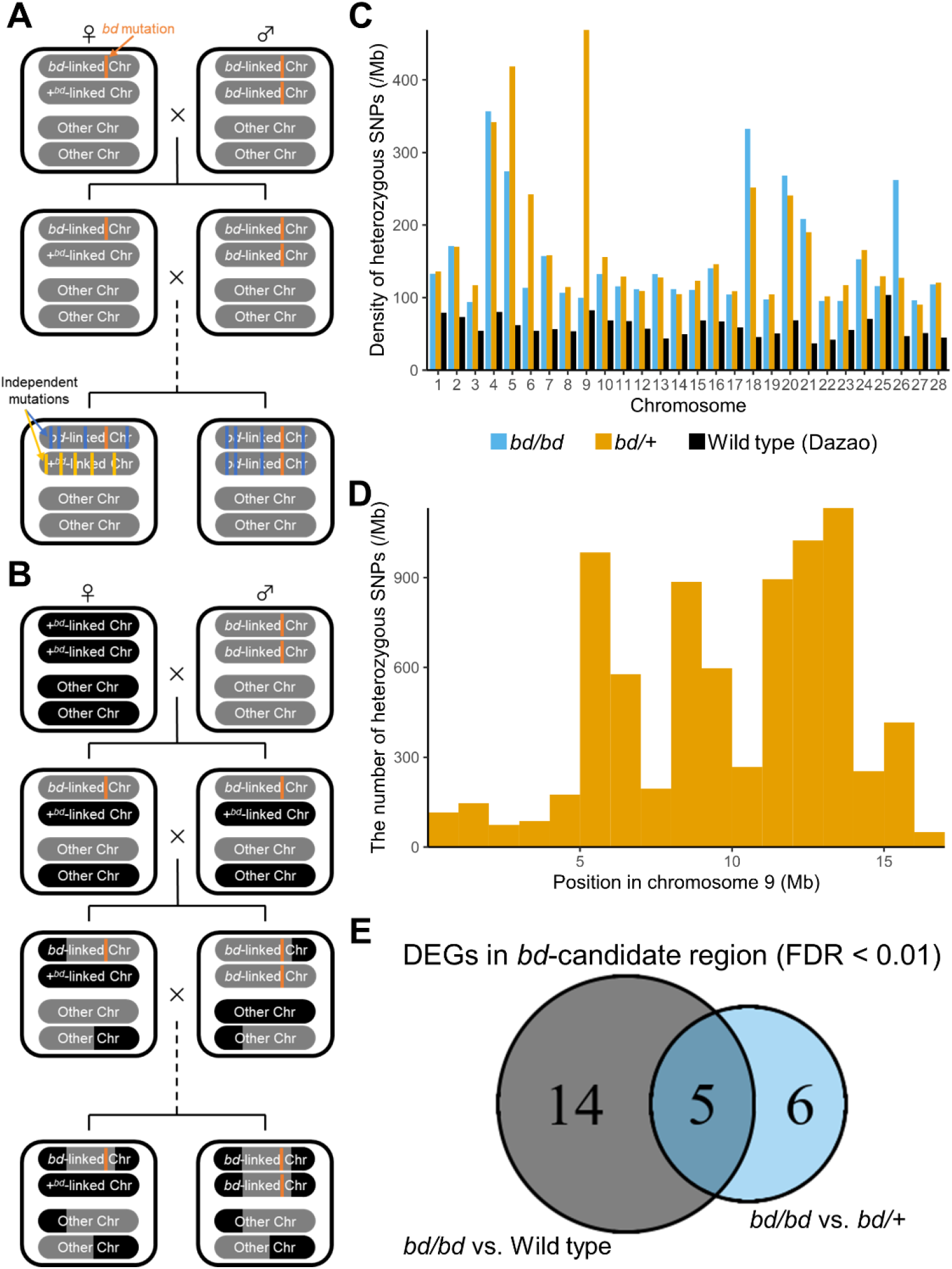
Prediction of *bd*-candidate genes from RNA-seq data. (A-B) The breeding history of the *bd* strain. If the *bd* strain has been maintained by sib-mating since its establishment, the genetic backgrounds of the *bd*-linked and *+^bd^*-linked chromosomes become heterozygous due to an accumulation of independent mutations (A). If the *bd* strain has been crossed with a strain other than Gunze-Ao and subsequently inbred, the origins of the *bd*-linked and *+^bd^*-linked chromosomes are different (B). (C) Density of heterozygous SNPs across all chromosomes in *bd/bd*, *bd/+*, and the wild-type Dazao strains. RNA-seq reads were mapped to the p50T reference genome (Kawamoto et al., 2019). (D) Number of heterozygous SNPs across chromosome 9 in *bd/+* individuals. (E) Number of genes within the *bd*-candidate region and differentially expressed between *bd/bd* and wild type (left) or between *bd/bd* and *bd/+* (right) analyses.

Wu et al. (2016) conducted an RNA-seq analysis using a mixture of total RNA extracted from the integument of four individuals from each of *bd/bd* mutant, *bd/+* mutant, and the wild-type Dazao. First, we reanalyzed their RNA-seq dataset to determine the chromosome location of the *bd* locus. We found a high number of heterozygous SNPs in chromosome 9, a result supported by a previous linkage analysis (Sasaki, 1941), in which the *bd/+* heterozygous individuals were compared with *bd/bd* and wild-type individuals (Figure 2C). Moreover, the number of heterozygous SNPs was higher in *bd/bd* mutants than in wild-type Dazao across all 28 chromosomes (Figure 2C), suggesting that *bd* strain is not sufficiently inbred. Thus, it is likely that the *bd* strain has been crossed with a strain other than Gunze-Ao and subsequently inbred, as shown in Figure 2B. Since the highest number of heterozygous SNPs was located in a region ranging from 5 to 16 Mb on chromosome 9 (Figure 2D), we hypothesized that the *bd* locus is located in this genomic region, hereafter the *bd*-candidate region. Next, we examined differentially expressed genes (DEGs; False Discovery Rate [FDR] < 0.01) located in the *bd*-candidate region. From this analysis, we detected several DEGs: 19 between wild-type and *bd/bd* strains, and 11 between *bd/+* and *bd/bd* strains (Figure 2E). Five genes, namely, *KWMTBOMO05086, KWMTBOMO05183, KWMTBOMO05216, KWMTBOMO05376*, and *KWMTBOMO05377*, were found to be differentially expressed in both analyses (Figure 2E, Table 1). The candidate gene responsible for the *bd* phenotype was expected to be either upregulated or downregulated in *bd/bd* mutants. We excluded the *KWMTBOMO05216* gene from the candidates since its expression level was higher in heterozygous than in both homozygous normal and homozygous mutants (*bd/+* > *bd/bd* > wild type). We then explored the closest relatives of each candidate in *D. melanogaster* using the BLASTP program in FlyBase (https://flybase.org/blast/) (Table 1). *KWMTBOMO05376* and *KWMTBOMO05377* genes are relative to *Cuticular protein 65Av* (*Cpr65Av*), which is predicted to be involved in chitin-based cuticle development (http://flybase.org/reports/FBgn0052405.html). The *KWMTBOMO05183* gene is relative to *CG9701*, which likely encodes a beta-glucosidase (https://flybase.org/reports/FBgn0036659.html). In consequence, there is no evidence that any of these three genes have a direct role in larval pigmentation or fertility. Moreover, and more importantly, we found that the *KWMTBOMO05086* gene is homologous to the *maternal gene required for meiosis* (*mamo*) gene which encodes a BTB-ZF transcription factor that is critical for female fertility in *D. melanogaster* (Mukai et al., 2007; Nakamura et al., 2019) (Figure S1). Given these results, we will refer to this homologous gene as “*B. mori mamo*” (*Bmmamo*) and propose that it is responsible for the *bd* phenotype.

**Table 1.**
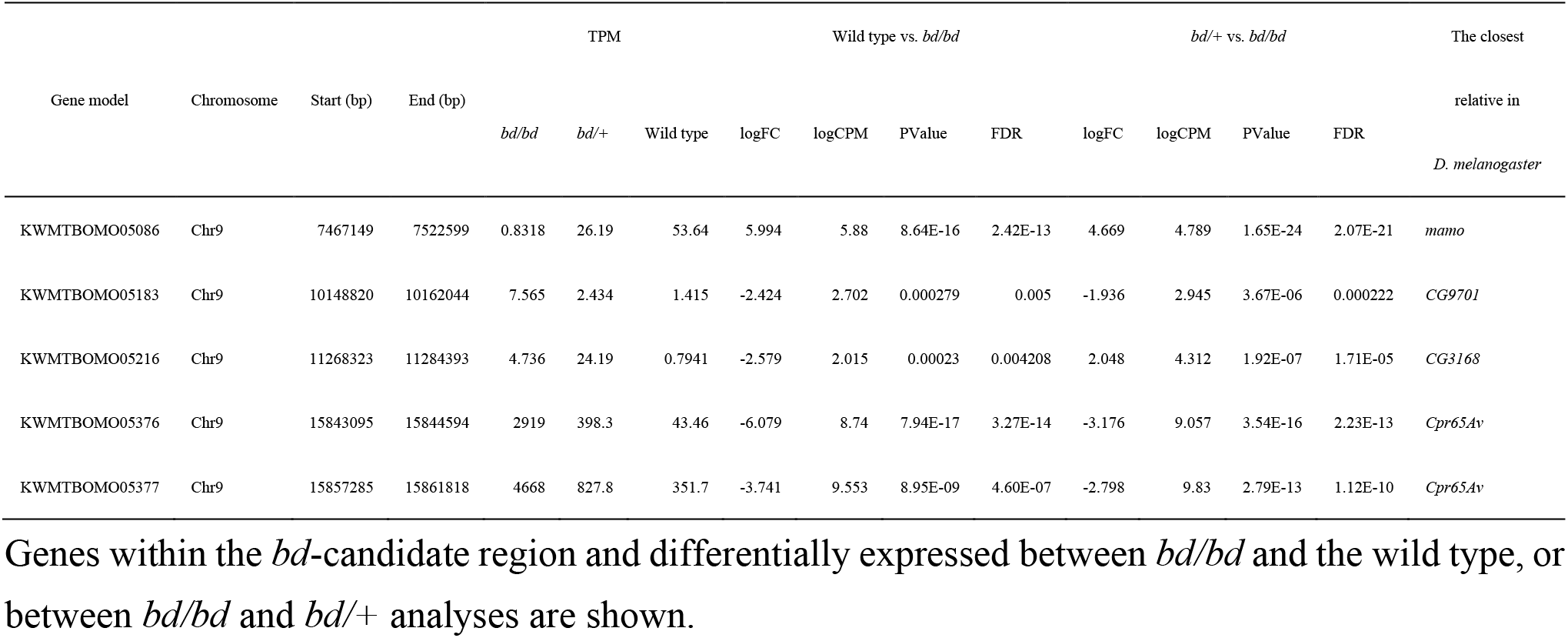
Candidate genes for the *bd* phenotype predicted using RNA-seq data.

### 3.2 Comparative analysis of the expression level of the *Bmmamo* gene

According to previous reports, most genes responsible for larval pigmentation in *B. mori* are highly expressed during or immediately before the molting period (Futahashi et al., 2010, 2008; Kondo et al., 2017; Wang et al., 2022; Yamaguchi et al., 2013). Therefore, we measured the expression level of the *Bmmamo* gene in the wild-type strain (p50T) at the fourth molting stage (Figure 3A). Developmental stage (C1, D1, E1, E2, and F) was determined on the basis of the spiracle index (Kiguchi and Agui, 1981). The expression level of the *Bmmamo* gene increased toward the fourth molting E1 stage, and then quickly decreased.

**Figure 3.**
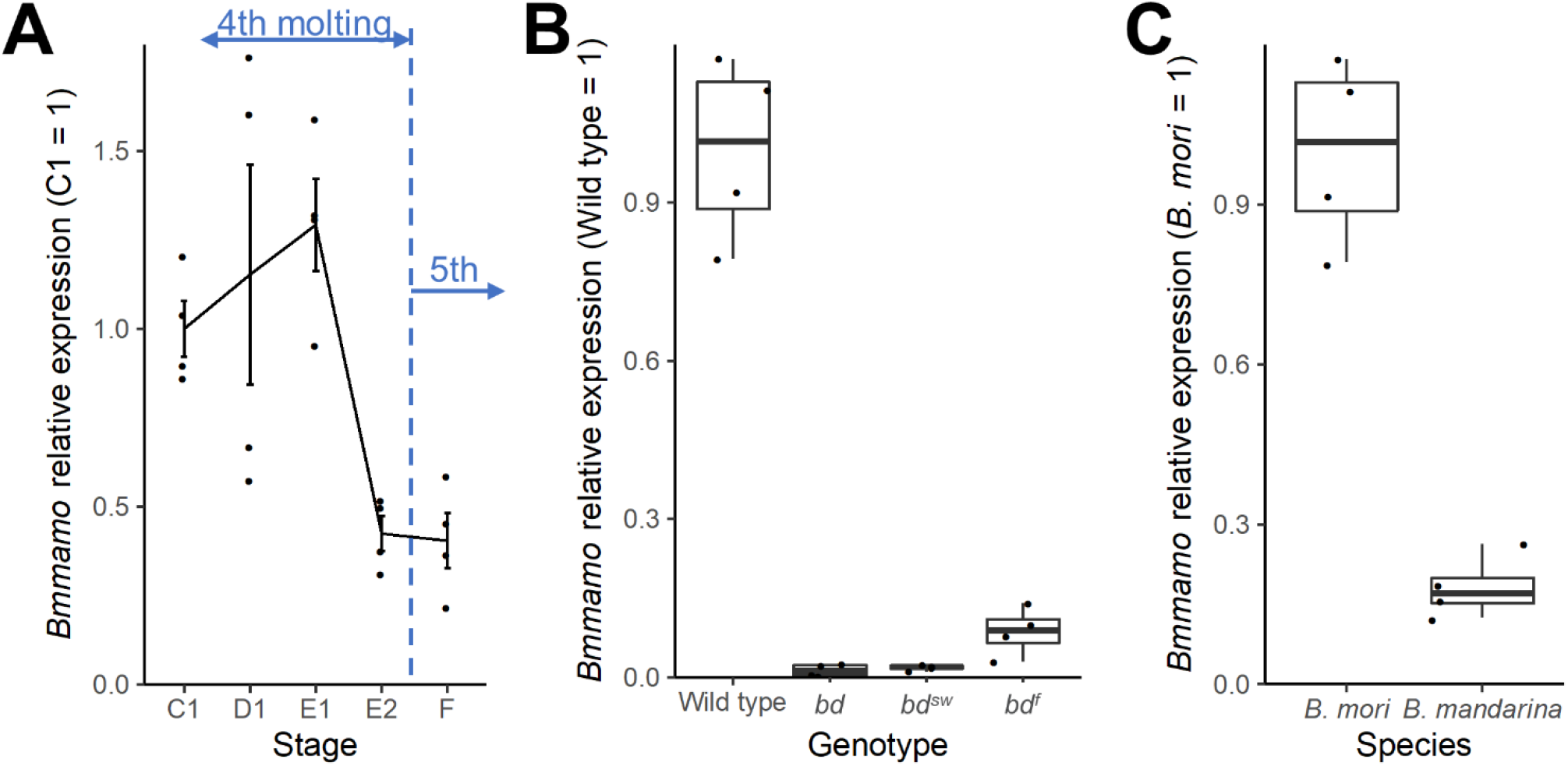
Analysis of *Bmmamo* expression. (A) Expression level of the *Bmmamo* gene in the integument of wild-type (p50T) larvae at the fourth molting period. The molting stage (C1, D1, E1, E2, and F) was determined on the basis of the spiracle index (Kiguchi and Agui, 1981). (B) Expression level of the *Bmmamo* gene in the larval integument of wild-type, *bd*, *bd^sw^*, and *bd^f^* larvae at the fourth molting E1 stage. (C) Expression level of the *Bmmamo* gene in the larval integument of *B. mori* and *B. mandarina* larvae at the fourth molting E1 stage. The expression level of the *Bmmamo* gene was normalized using the housekeeping gene *ribosomal protein 49* (*rp49*) as reference gene.

We compared *Bmmamo* expression in the larval integument of wild-type (p50T) and *bd* strains at the E1 stage (Figure 3B). The expression of the *Bmmamo* gene was lower in *bd*, *bd^sw^*, and *bd^f^* mutants than in the wild type. We then cloned the coding sequence of the *Bmmamo* gene only in wild-type and *bd^f^* strains since its expression level was undetectable in *bd* and *bd^sw^* mutants (Figure S2). The BmMamo protein of wild-type (p50T) and *bd^f^* strains showed three amino acid substitutions, namely, T611S, H695R, and P742L. The former two substitutions were observed in 13 and 7 of the other 18 +*^bd^* strains, respectively, but the latter substitution was not found in any of the +*^bd^* individuals according to the hypothetically reconstructed genome data (Kawamoto et al., 2022) (Table S2). According to InterProScan (Paysan-Lafosse et al., 2023), P742L is located within the zinc finger C2H2 domain (Figure S3A). In addition, the level of the *Bmmamo* gene at the E1 stage was higher in *B. mori* than in *B. mandarina* larvae (Figure 3C), suggesting that it may be involved in larval color differences between two species.

### 3.3 CRISPR/Cas9-mediated knockout of the *Bmmamo* gene

We designed a crRNA targeting the fifth exon that is common to all isoforms of the *Bmmamo* gene (Yokoi et al., 2021; https://kaikobase.dna.affrc.go.jp/KAIKObase/gbrowse/chromosome_2_3/?name=id:336414;dbid=chromosome_2_3:database) and performed CRISPR/Cas9-mediated mutagenesis (Figure 4A). From this technique, we obtained 19 of 23 fifth instar larvae that exhibited mosaic-mottled black patches on the normal white integument (Table S3, Figure S4). We then crossed these G_0_ moths to wild-type moths and obtained G_1_ individuals. The sib-cross of G_1_ individuals allowed us to obtain three different *Bmmamo* strains with homozygous mutations at the target site, namely, *Bmmamo^Δ4^*, *Bmmamo^Δ15A^*, and *Bmmamo^Δ15B^* (Figure 4A). In further studies, we used the *Bmmamo^Δ4^* strain since it harbored a frameshift mutation with a premature stop codon (Figure 4A). *Bmmamo^Δ4^* larvae had grayish-black color (Figure 4B), and *Bmmamo^Δ4^* adults were viable and developed abnormally small wings (Figure 4C), a similar phenotype as observed in *bd* mutants (Sasaki, 1941). In addition, *Bmmamo^Δ4^* females laid fewer eggs than the wild type, most of them were colorless (Figure 4D) and none hatched (Table S4). Since normal eggs start to take color from around 40 hours after oviposition, egg development is likely stopped at an early stage or even before fertilization. This observation supports the idea that a failure of the micropylar apparatus caused sterility in *bd* and *bd^sw^* females (Kawaguchi et al., 2006; Sakanoshita, 1970). Moreover, the complementation cross between *Bmmamo^Δ4/+^* and *bd/+* strains produced approximately a quarter of larvae with grayish-black color in a similar way to the observed in *bd* mutants (Figure 4B). These results indicate that mutations in the *Bmmamo* gene are responsible for the *bd* phenotypes, i.e., grayish-black pigmentation in larvae and sterility in females. Additionally, we observed that *Bmmamo^Δ4^* knockout mutant males cannot mate themselves (see Supplemental movie for more detail) and require hand-paring, similarly as previously described in *bd* males (Sasaki, 1941).

**Figure 4.**
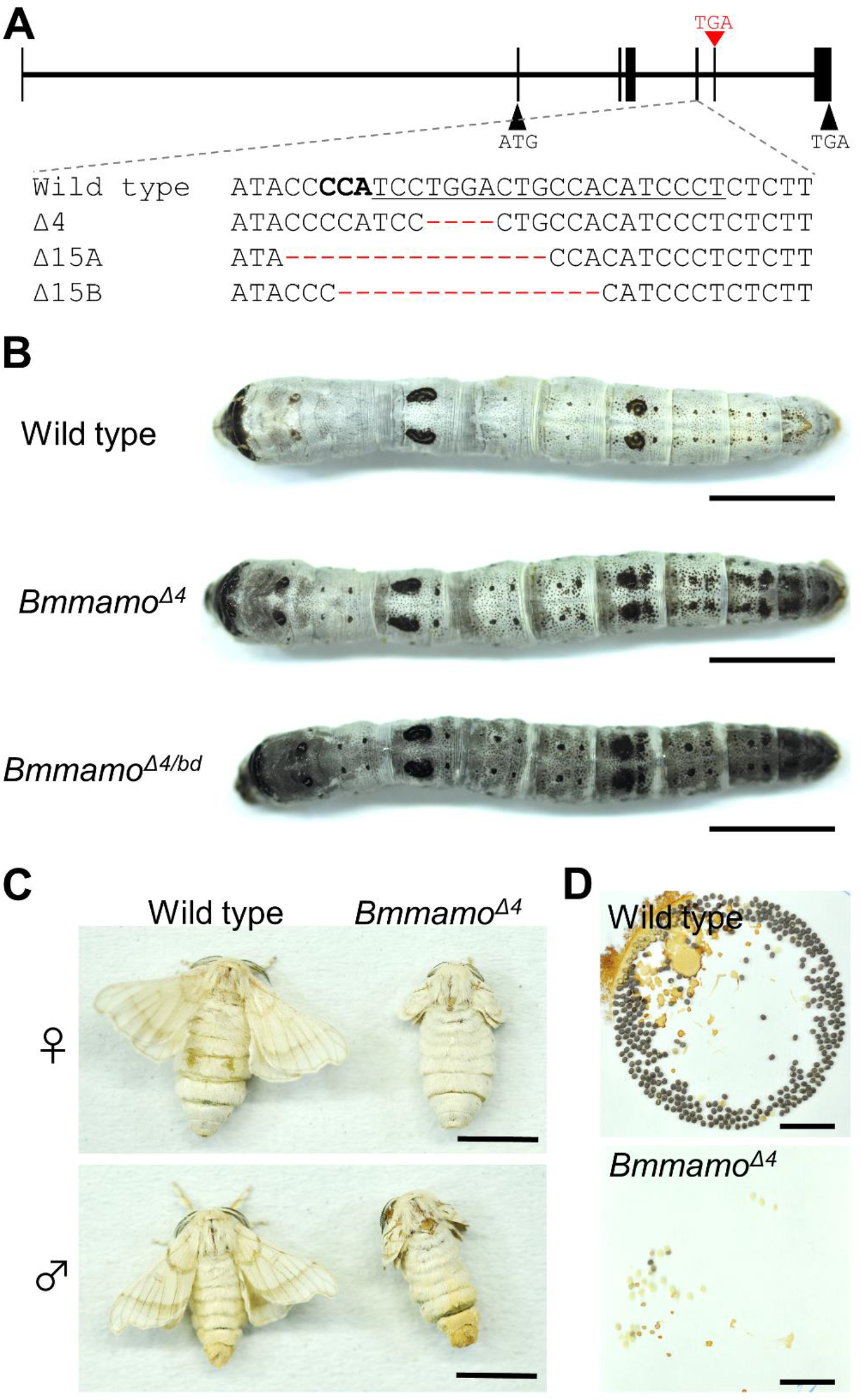
CRISPR/Cas9-mediated knockout of the *Bmmamo* gene. (A) Exon structure of the *Bmmamo* gene and sequence alignment surrounding the crRNA target site in wild-type (p50T) and *Bmmamo* knockout strains. The target sequence is underlined, and the protospacer adjacent motif site is shown in bold letters. A red triangle indicates the premature stop codon of the *Bmmamo^Δ4^*mutation. (B) Representative wild-type, *Bmmamo* knockout (*Bmmamo^Δ4^*), and *Bmmamo^Δ4/bd^* larvae obtained from complementation cross between *Bmmamo^Δ4/+^* and *bd/+* mutants. *Bmmamo^Δ4/bd^* larvae exhibited a *bd*-like body color. (C) Phenotypes of wild-type adult female (top left), *Bmmamo^Δ4^* adult female (top right), wild-type adult male (bottom left), and *Bmmamo^Δ4^* adult male (bottom right). (D) Eggs at five days after oviposition obtained by crossing a wild-type female to a wild-type male (top) and a *Bmmamo^Δ4^* female to a wild-type male (bottom). Scale bars: 1 cm.

### 3.4 Analysis of expression of melanin synthesis genes in *Bmmamo* mutants

The previous RNA-seq analysis detected DEGs that may be involved in the larval pigmentation of *bd* mutants, including a melanin synthesis gene *laccase2* (Wu et al., 2016). However, it is worth noting that other melanin synthesis genes such as *tyrosine hydroxylase* (*TH*), *yellow-y*, *ebony*, and *DOPA decarboxylase* (*DDC*) were not differentially expressed. We propose that this result is due to the fact that the total RNA was extracted from the larval integument at the early molting stage. In addition, previous analysis was performed using strains with different genetic backgrounds, which may bias the result. Since *Bmmamo^Δ4^*individuals originated using the p50T strain, it is possible to eliminate genetic background biases by comparing both strains.

We performed a qPCR analysis targeting eight melanin synthesis genes using total RNA extracted from the larval integument of wild-type (p50T) and *Bmmamo^Δ4^* knockout strains at E1 and E2 stages, during and slightly after the peak in the expression of the *Bmmamo* gene (Figure 3A). The expression of *DDC*, *tan*, and *yellow-y* genes was significantly higher in *Bmmamo^Δ4^* knockout mutants than in the wild-type strain either at E1 or E2 stages (Figure 5). Although statistically non-significant, the expression levels of *TH* and *laccase2* were also higher in *Bmmamo^Δ4^* knockout mutants than in the wild type (Figure 5). These results showed that the normal BmMamo protein directly or indirectly suppresses the expression of melanin synthesis genes, especially in genes involved in the synthesis of black melanin pigments (i.e., Dopa melanin and dopamine melanin pigments) (Figure 6).

**Figure 5.**
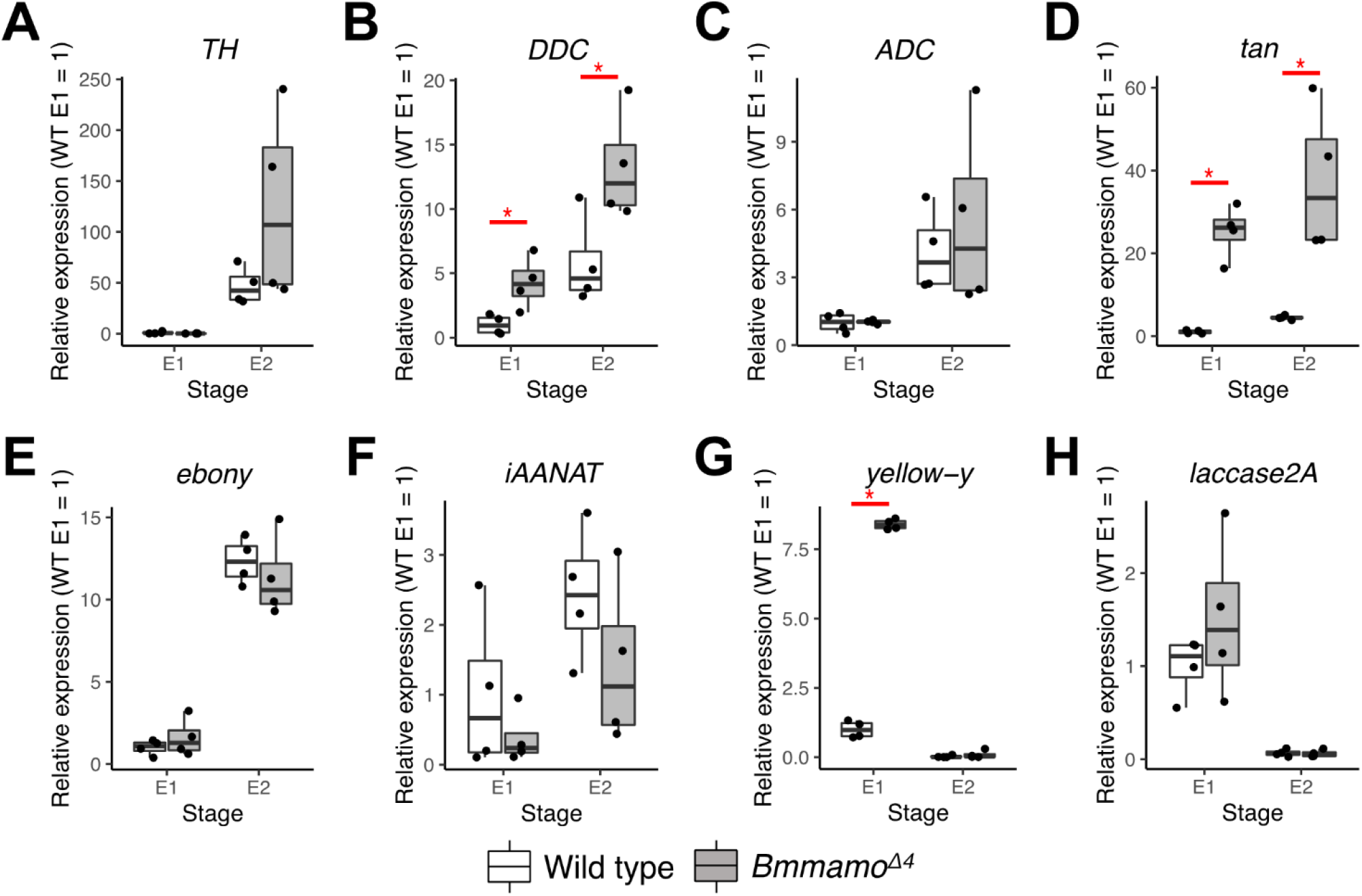
Expression levels of melanin synthesis genes in the integument of wild-type and *Bmmamo^Δ4^*strains at fourth molting E1 and E2 stages. Expression levels of *TH* (A), *DDC* (B), *ADC* (C), *tan* (D), *yellow-y* (E), and *laccase2* (F) genes were normalized using *rp49* as reference gene. The *B. mori* genome harbors two *laccase2* genes, *laccase2A* and *laccase2B*, but only the expression level of *laccase2A* was measured since it is believed to have a major role in cuticle formation (Yatsu and Asano, 2009). Asterisks indicate statistically significant values in Welch’s *t* test (p < 0.05).

**Figure 6.**
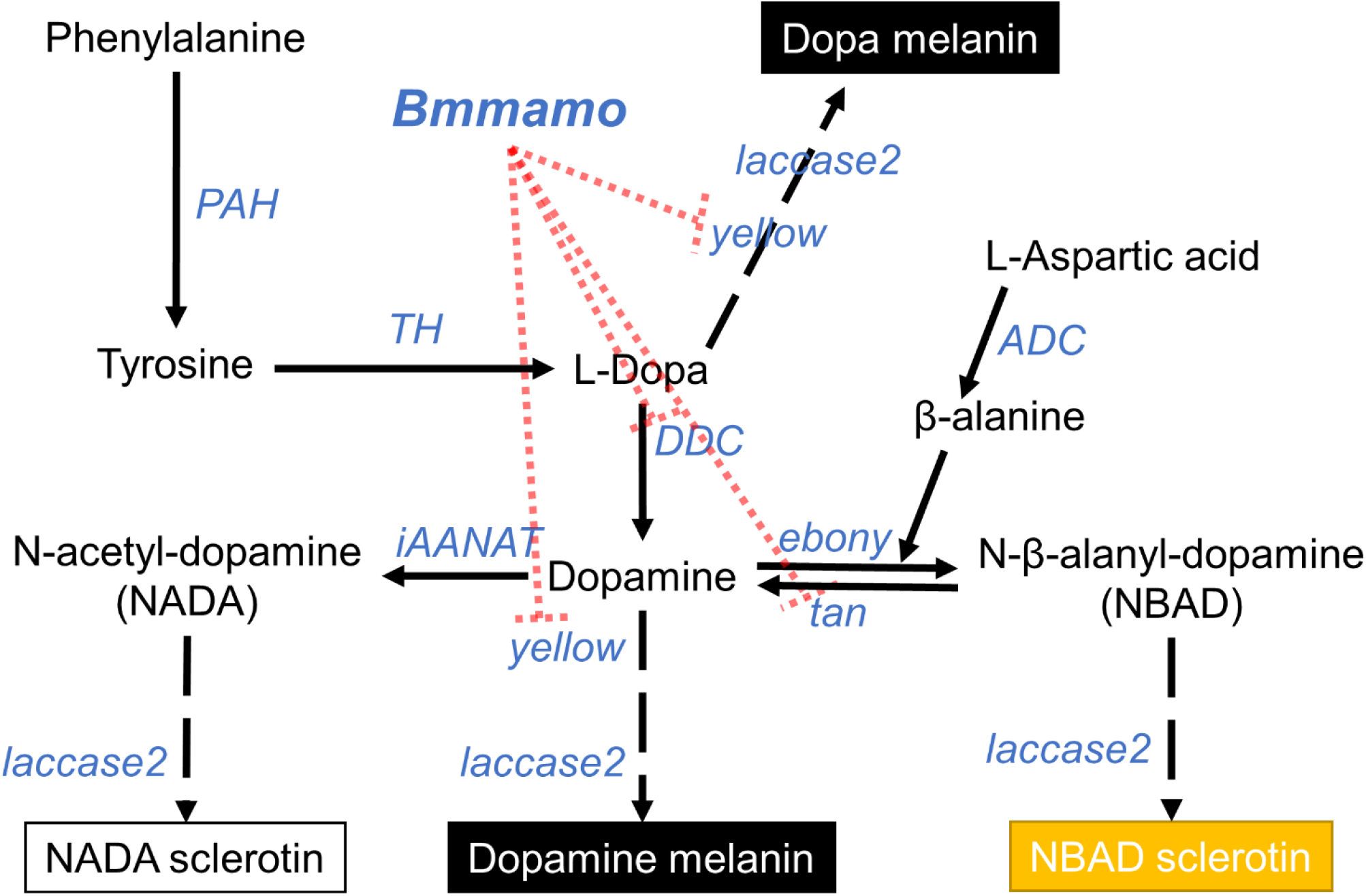
Schematic models of the regulatory network involving *Bmmamo* and melanin synthesis genes. The melanin biosynthesis pathway in *Bombyx* larvae has been adapted from Futahashi et al. (2010, 2008), Zhan et al. (2010), and Yoda et al. (2014). Because it remains unknown whether BmMamo protein directly or indirectly suppresses the expression of melanin synthesis genes, the interactions are indicated by dotted lines.

## 4 Discussion

### 4.1 Functional conservation and divergence of homologous *mamo* genes

To our knowledge, *D. melanogaster* is the only insect other than *B. mori* in which the function of the *mamo* gene has been analyzed. In *D. melanogaster*, homozygous *mamo* mutants are lethal at the third instar stage (Mukai et al., 2007). The FLP/FRT/DFS system (Chou and Perrimon, 1992) was used to obtain *mamo* homozygous germline clones and embryos showed lethality at late embryonic developmental stages (stages 16 and 17) (Mukai et al., 2007). It has been shown that the maternal Mamo protein regulates the expression of the *vasa* gene during embryogenesis (Nakamura et al., 2019), an observation that may explain the lethality of embryos. On the other hand, in *B. mori*, we observed that *Bmmamo* knockout mutants were viable and developed into adult moths (Figure 4BC). *Bmmamo* knockout females laid unhatched eggs due to deficiencies in fertilization (Figure 4D, Table S4), which is likely due to the failure of the micropylar apparatus (Kawaguchi et al., 2006; Sakanoshita, 1970).

In this study, we observed that *Bmmamo* loss-of-function results in an increase in larval melanization. However, in *D. melanogaster*, a relationship between the *mamo* gene and the melanin synthesis pathway has not been reported yet (https://flybase.org/reports/FBgn0267033). Similarly, some other transcription factor genes involved in larval pigmentation in *B. mori* have not been reported to affect coloration in *D. melanogaster*. For example, transcription factor genes *apt-like* and *SoxD* are known to affect the pigmentation pattern of *B. mori* larvae (Wang et al., 2022; Yoda et al., 2014). However, genes relative to *apt-like* and *SoxD* in *D. melanogaster*, *apt* and *Sox102F*, respectively, have not been reported to affect coloration (https://flybase.org/reports/FBgn0039938; https://flybase.org/reports/FBgn0015903; Note that *apt-like* and *apt* share sequence similarity but are not orthologous). We do not know whether these transcription factors acquired new functions in Lepidoptera or lost their functions in Diptera, thereby further studies at the family taxonomic level are needed to elucidate the role of these genes on insect pigmentation.

Taken together, our results support that *mamo* and *Bmmamo* are orthologous genes with different functions. The *mamo* gene is a member of the BTB-ZF protein family. To date, 16, 13, 13, 3, 58, and 63 BTB-ZF proteins have been characterized in *B. mori*, *Danaus plexippus*, *D. melanogaster*, *Caenorhabditis elegans*, *Mus musculus*, and *Homo sapiens*, respectively (Wu et al., 2019). This high variation in the number of BTB-ZF proteins in different species suggests that BTB-ZF genes may have undergone either duplication and/or loss events during the evolution of each lineage.

### 4.2 Evaluation of pleiotropic effects of the *Bmmamo* gene

We found that the *Bmmamo* gene is associated with both larval pigmentation and female fertility. In *D. melanogaster*, *mamo* is involved in neural development and differentiation (Lai et al., 2022; Liu et al., 2019). In *B. mori*, it has been reported that the *Bmmamo* gene is highly expressed in the adult brain (https://silkbase.ab.a.u-tokyo.ac.jp/cgi-bin/entryview_gm2.cgi?clone_name=%20KWMTBOMO05086). It is difficult to examine behavioral effects in *Bmmamo* mutants since knockout adults have abnormal small wings (Figure 4C), but *Bmmamo* knockout males exhibit abnormal mating behavior (Supplemental movie). Taken together, these observations suggest that the *Bmmamo* gene may also play a role in neural development.

The phenomenon by which a single locus affects two or more phenotypic traits is called pleiotropy. According to their effects on the phenotype, pleiotropy can be classified into two types: 1) type I pleiotropy, conferred by multiple molecular functions of a gene, and 2) type II pleiotropy, associated with multiple morphological and physiological consequences of a single molecular function (Wagner and Zhang, 2011). There are a few examples of type I pleiotropy (Uggè et al., 2022), and previous findings suggest that pleiotropic effects are mostly of type II (He and Zhang, 2006; Wagner and Zhang, 2011). In *D. melanogaster*, the *mamo* gene yields seven splice isoforms that encode four different proteins. Only one Mamo isoform is involved in the generation and maintenance of α’/β’ and γ neurons in the mushroom body (Lai et al., 2022; Liu et al., 2019), suggesting that the other isoforms are involved in different functions. Nakamura et al. (2019) showed that the maternally derived Mamo protein is processed into shorter derivatives without BTB domain. Both full-length and truncated Mamo proteins are likely involved in oogenesis and embryogenesis, but the localization and function of these isoforms are different (Nakamura et al., 2019). In *B. mori*, transcriptomic analysis indicates that the *Bmmamo* mRNA is spliced into several transcripts coding for multiple protein isoforms (Yokoi et al., 2021; https://kaikobase.dna.affrc.go.jp/KAIKObase/gbrowse/chromosome_2_3/?name=id:336414;dbid=chromosome_2_3:database). Taken together, the *Bmmamo* gene is suggested to be subject to type I pleiotropic effects, i.e., multiple protein isoforms with distinct molecular functions generated by a single gene. Our next goal is to reveal the functional differentiation of these BmMamo isoforms. Long read RNA-seq at different time points in multiple tissues can be used to create an atlas of *Bmmamo* gene isoform expression. In this study, we performed CRISPR/Cas9-mediated knockout targeting the fifth exon of the *Bmmamo* (Figure 4A). Functional differences in BmMamo isoforms can be assessed by CRISPR/Cas9-mediated mutagenesis targeting other exons and comparing the phenotypes of the mutant strains.

### 4.3 Domestication-related pigmentation patterns in *B. mori*

Genes selected during domestication and associated with color variation between *B. mori* and *B. mandarina* (e.g., *apt-like*, *SoxD*, and *ADC* genes) have recently been identified by investigating *B. mori* mutants (Dai et al., 2015; Wang et al., 2022; Yoda et al., 2014). In addition, previous comparative studies exploring the degree of sequence divergence between *B. mori* and *B. mandarina* suggest that melanin synthesis genes such as *TH*, *insect arylalkylamine N acetyltransferase* (*iAANAT*), and *ADC* may have experienced artificial selection during silkworm domestication (Tong et al., 2022; Xiang et al., 2018; Yu et al., 2011). These observations support the idea that domestication-related pigmentation changes in *B. mori* is a polygenic and complex trait.

Importantly, a recent sequence comparative analysis suggested that the *Bmmamo* gene has been selected during silkworm domestication (Tong et al., 2022). The BmMamo protein suppresses the expression of melanin synthesis genes (Figure 5), and its expression in the larval integument is higher in *B. mori* than in *B. mandarina* (Figure 3C). Taken together, these observations indicate that the *Bmmamo* may be involved in domestication-associated color changes of *B. mori*.

The molecular mechanism by which *Bmmamo* contributes to pigmentation variation remains to be elucidated. For example, the *apt-like* gene associated with differential pigmentation of *B. mori* and *B. mandarina* larvae (Tomihara et al., 2022) is expressed in the larval integument, and its level reaches an earlier peak than that of the *Bmmamo* gene at the fourth molting stage (Figure S5). In consequence, we propose that the *Bmmamo* gene may be regulated by the *apt-like* gene. However, the expression level of the e*bony* gene, which is under the control of the *apt-like* gene (Futahashi et al., 2008), was not induced by *Bmmamo* knockouts (Figure 5). Further studies investigating the molecular relationship between *Bmmamo* and *apt-like* genes are needed to unravel the underlying genetic basis of the larval pigmentation trait associated with selection during silkworm domestication.

### 4.4 The phenotypic variations observed in *bd* and *Bmmamo* knockout mutants

The larval integument of the *bd* mutant (k40 strain) is darker compared to that of the *Bmmamo^Δ4^* knockout mutant (p50T strain) (Figure S6). The integument of *bd/+* larvae is also darker than that of *Bmmamo^Δ4/+^* larvae (Figure S6). Among *plain* (*p*) alleles, *+^p^* (or *P^3^*) larvae have eye spots, crescent spots, and star spots, whereas *p* larvae do not (Yoda et al., 2014). Previous studies found that *bd* larvae on *+^p^* backgrounds have a darker color compared to *p* backgrounds (Aruga et al., 1954; Sasaki, 1956). In addition, double mutants of *bd* and other mutations responsible for larval markings, namely *Striped* (*P^S^*), *U*, *Zebra* (*Ze*), and *Multi lunar* (*L*), are darker than *bd* larvae on *+^p^*background (Aruga et al., 1954; Sasaki, 1956). These results indicate that genetic backgrounds affect the larval color of *bd* mutants. Therefore, the difference in larval color between the *bd* mutant and the *Bmmamo^Δ4^*knockout mutant can be explained by their different genetic backgrounds. This can also explain the color difference between *Bmmamo^Δ4/bd^* and *Bmmamo^Δ4^*larvae (Figure 4B). However, the truncated Bmmamo protein containing only the BTB domain produced in the *Bmmamo^Δ4^* knockout mutant may retain some activity in repressing melanin synthesis genes (Figure S3B). Similarly, the *bd* and *bd^sw^* are thought to be null mutants because the expression of the *Bmmamo* gene was not detected (Figure 3B, Figure S2), but they have different larval colors (Figure 1D), which can be explained by their different genetic backgrounds.

Interestingly, while *bd^f^* larvae exhibit a grayish-black color, their females are fertile (Shimizu and Matsuno, 1978). We found that an amino acid substitution P742L in the BmMamo is specific to the *bd^f^*mutant strain (Table S2). It is possible that P742L is involved in larval pigmentation but does not affect female fertility. Alternatively, another hypothesis can be proposed. *Bmmamo^Δ4/+^* larvae are slightly darker than wild-type larvae (Figure S6), demonstrating that *Bmmamo* dosage is critical for larval pigmentation. The expression level of the *Bmmamo* gene in the larval integument of the *bd^f^* mutant is lower than that of the wild type (Figure 3B). Thus, it is also possible that in the *bd^f^* mutant, a mutation in the promoter or cis-regulatory region of the *Bmmamo* gene leads to its decreased expression in the integument and increased melanization, although the protein is still functional in the ovary. To clarify our hypotheses, it is required to introduce the P742L mutation into the wild-type strain using TALEN or CRISPR/Cas9-mediated knock-in approaches (Heryanto et al., 2022; Takasu et al., 2016). In addition, whole genome sequencing of the *bd^f^* strain may reveal the potential mutation responsible for *bd^f^*phenotypes in non-coding regions surrounding the *Bmmamo* gene.

## Data availability

The nucleotide sequences of *Bmmamo* were submitted to the DDBJ under accession numbers of LC760476 (p50T) and LC760477 (*bd^f^*).

## Acknowledgements

We are grateful to Susumu Katsuma for useful comments. We thank the Institute for Sustainable Agro-ecosystem Services at the University of Tokyo for facilitating the mulberry cultivation and the Biotron Facility at the University of Tokyo for rearing the silkworms. This work was supported by JSPS KAKENHI Grant Number JP20J22954 to KT. This work was also supported by the grant from the Ministry of Agriculture, Forestry and Fisheries of Japan (Research Project for Sericultural Revolution) to TK.

## Author contributions

KT designed the study and TK supervised it. KT conducted most of the experiments and analyses. KT wrote the manuscript with input from TK. All authors approved the final version of the manuscript and agree to be accountable for all aspects of the work.

## Notes

### Competing Interest Statement

The authors have declared no competing interest.

### Summary of Updates

This version of the manuscript has been revised following the editor's and reviewers' comments.

