## Supplemental figures for "Disruption of a BTB-ZF transcription factor causes female sterility and melanization in the larval body of the silkworm, *Bombyx mori*"

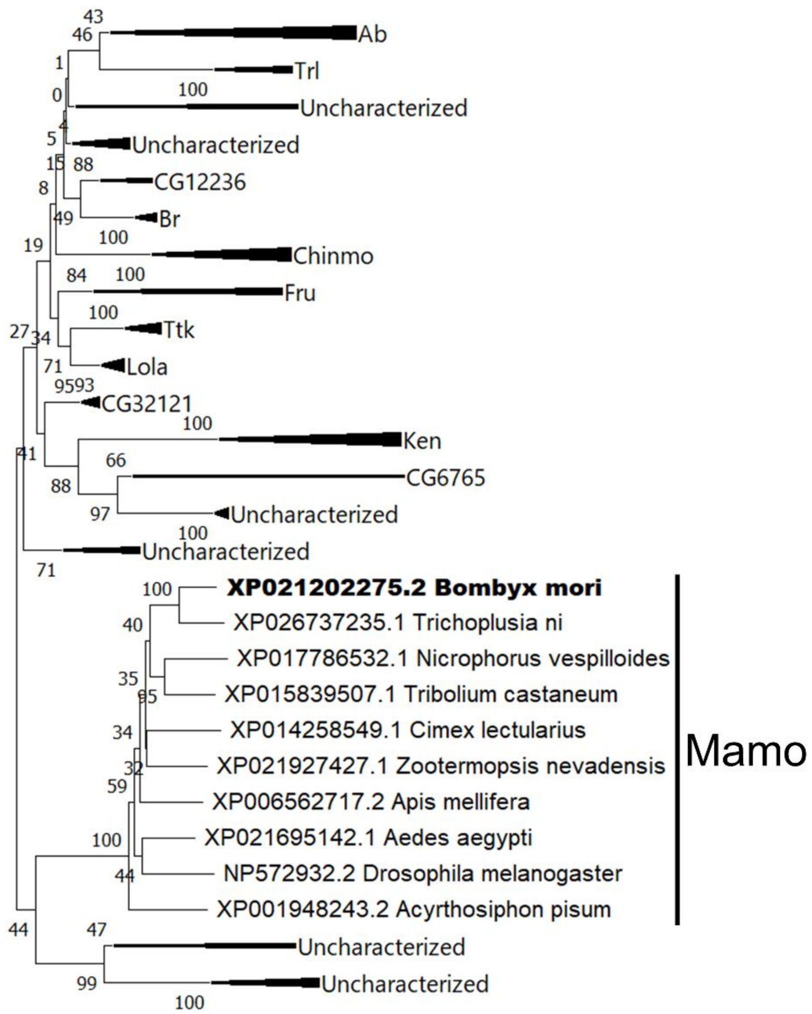


**Figure S1.** Phylogenetic analysis of BTB-ZF proteins in 12 insect species (*B. mori*, *T. ni*, *D. melanogaster*, *M. domestica, A. aegypti, A. mellifera*, *B. terrestris*, *T. castaneum*, *N. vespilloides*, *A. pisum*, *C. lectularius*, and *Z. nevadensis*). Accession numbers are given with species names. BmMamo is shown in bold letters. Collapsed clades are shown as triangles in protein groups other than Mamo homologs. The percentage of replicate trees in which the associated taxa clustered together in the bootstrap test (1000 replicates) are shown next to the branches. Ab; Abrupt, Trl; Trithorax-like, Br; Broad, Chinmo; Chronologically inappropriate morphogenesis, Fru; Fruitless, Ttk; Tramtrack, Lola; Longitudinals lacking, Ken; Ken and barbie.

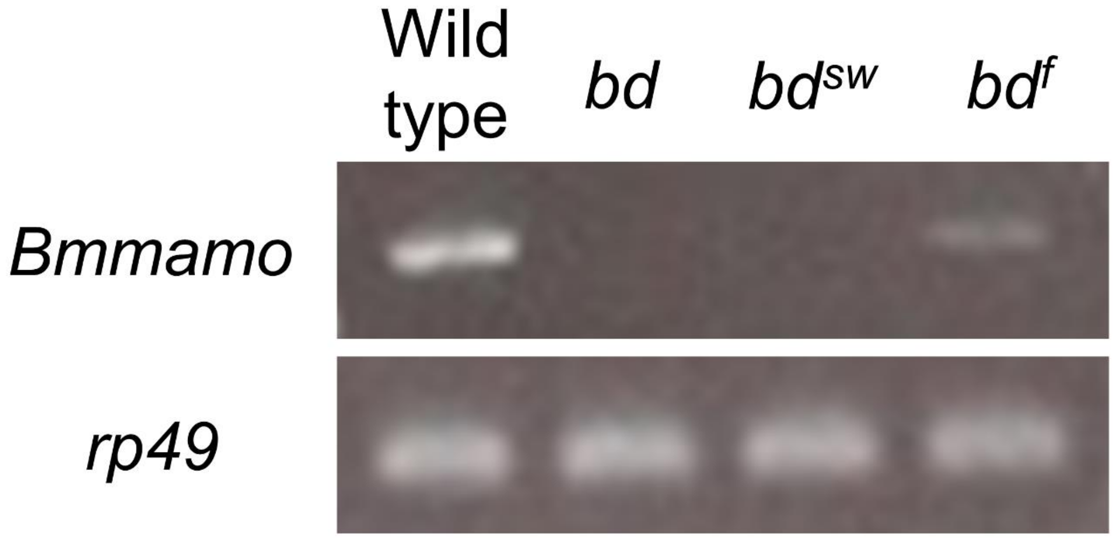


**Figure S2.** Gel electrophoresis of RT-PCR products using cDNA from integuments of wild-type and *bd* strains. The *Bmmamo* coding sequence was amplified using primers as shown in Table S1. The *rp49* gene was used as a reference control.

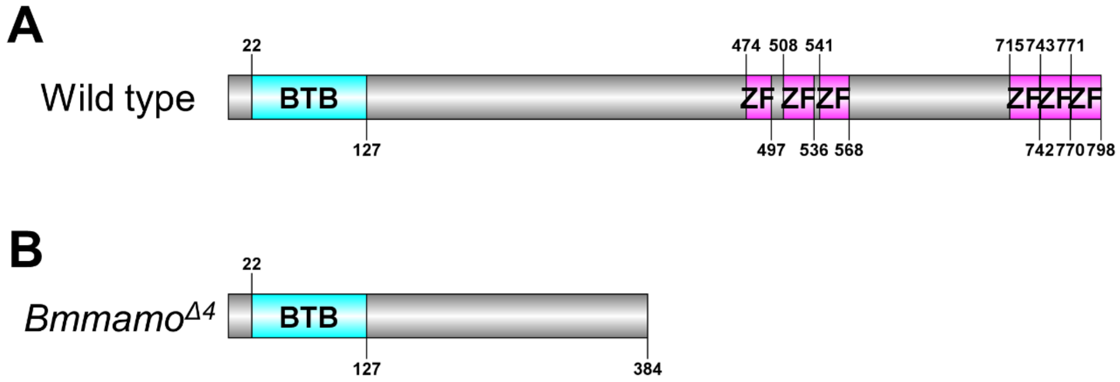


**Figure S3.** BmMamo protein structure of wild-type (A) and *Bmmamo*$^{\Delta4}$ (B) strains. Domains were predicted by InterProScan (Paysan-Lafosse et al., 2023). BTB: BTB/POZ domain (https://www.ebi.ac.uk/interpro/entry/InterPro/IPR000210/), ZF: Zinc finger C2H2-type domain (http://www.ebi.ac.uk/interpro/entry/InterPro/IPR013087/).

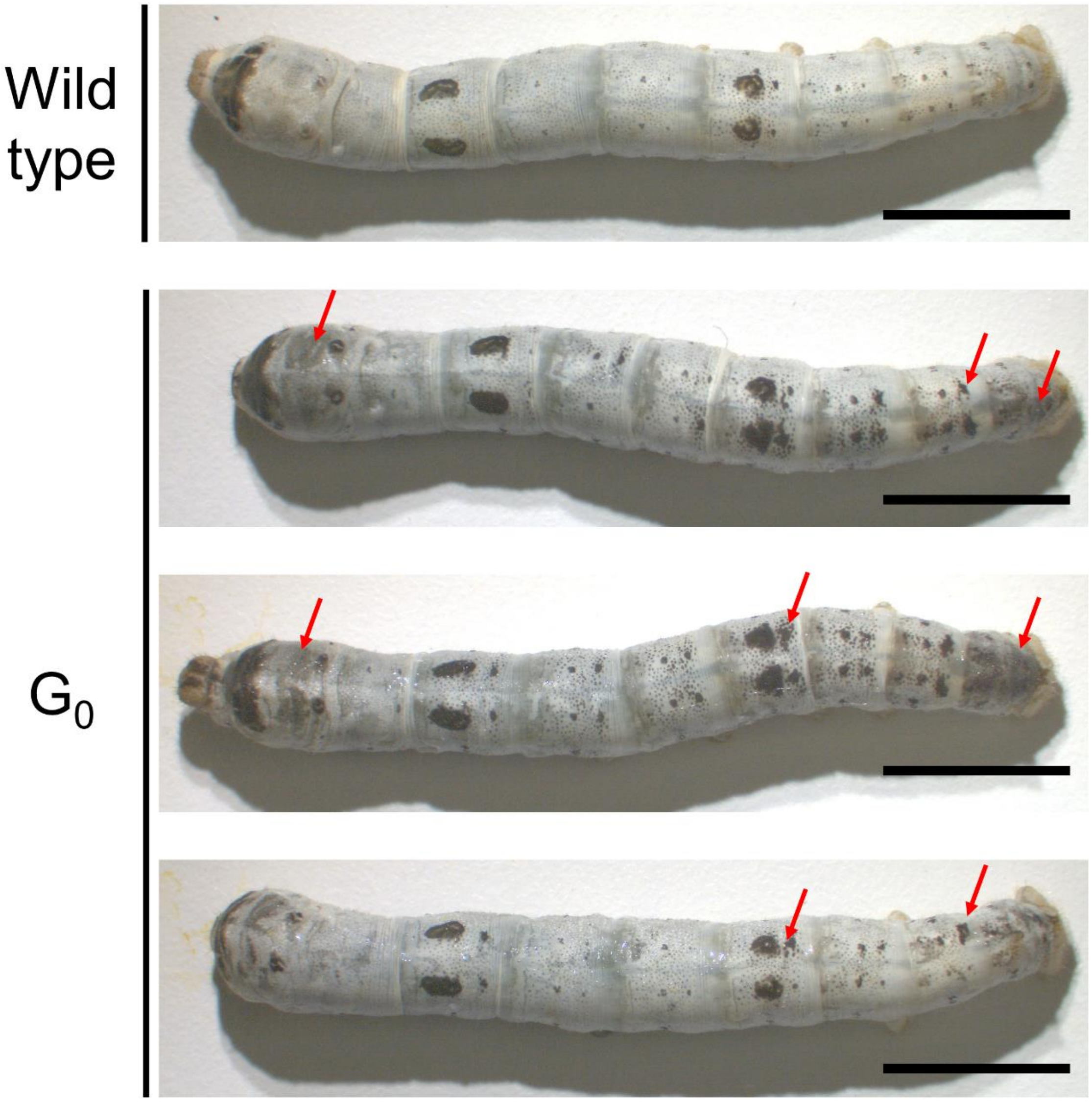


**Figure S4.** Larval phenotypes of *Bmmamo* knockout $G_0$ mutants. Most of the $G_0$ larvae exhibited mosaic-mottled black patches (pointed by red arrows) in the white integument.

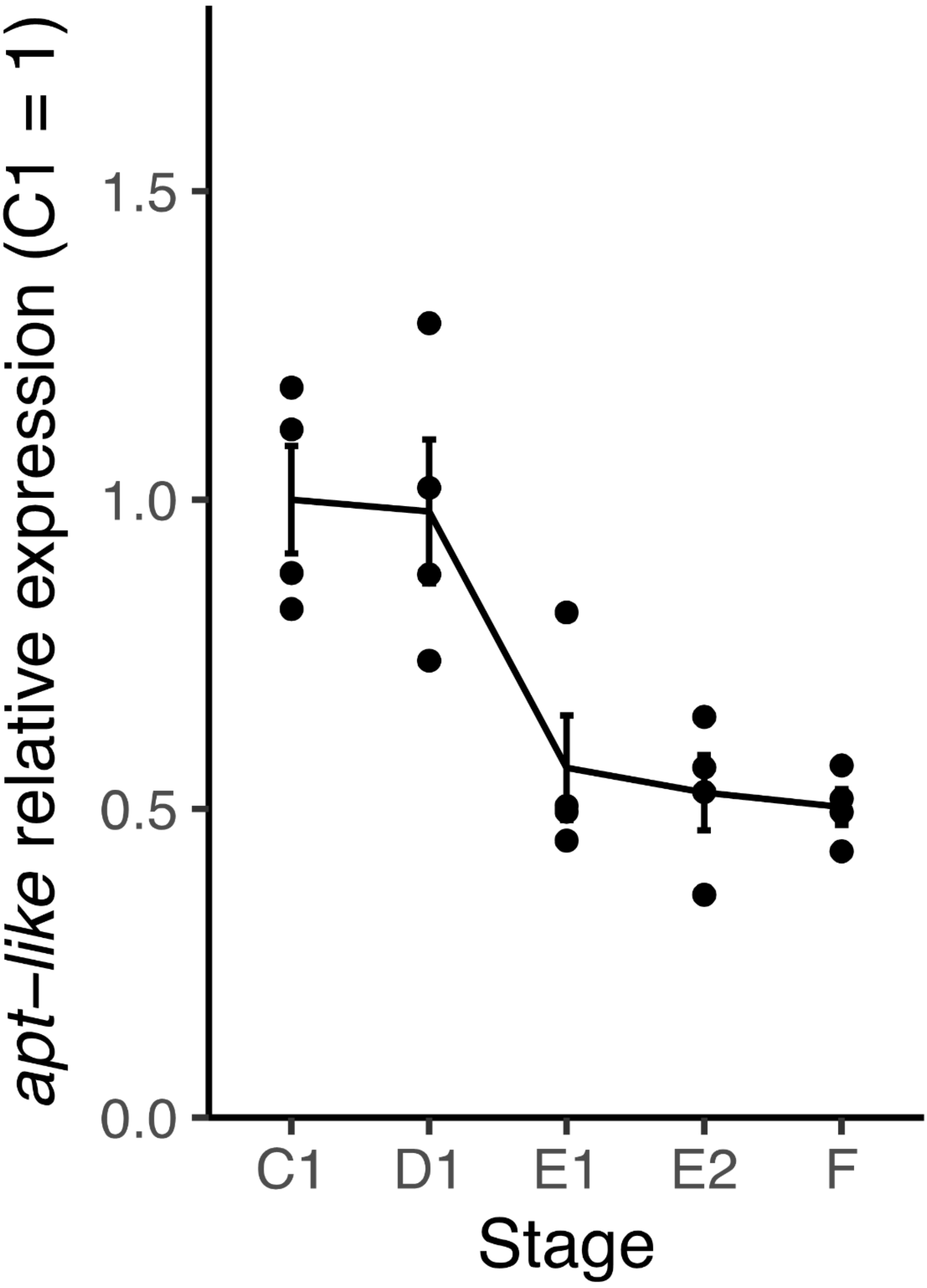


**Figure S5.** Expression level of the *apt-like* gene in the integument of wild-type p50T larvae at the fourth molting period. The expression level of the *apt-like* gene was normalized using *rp49* as reference gene.

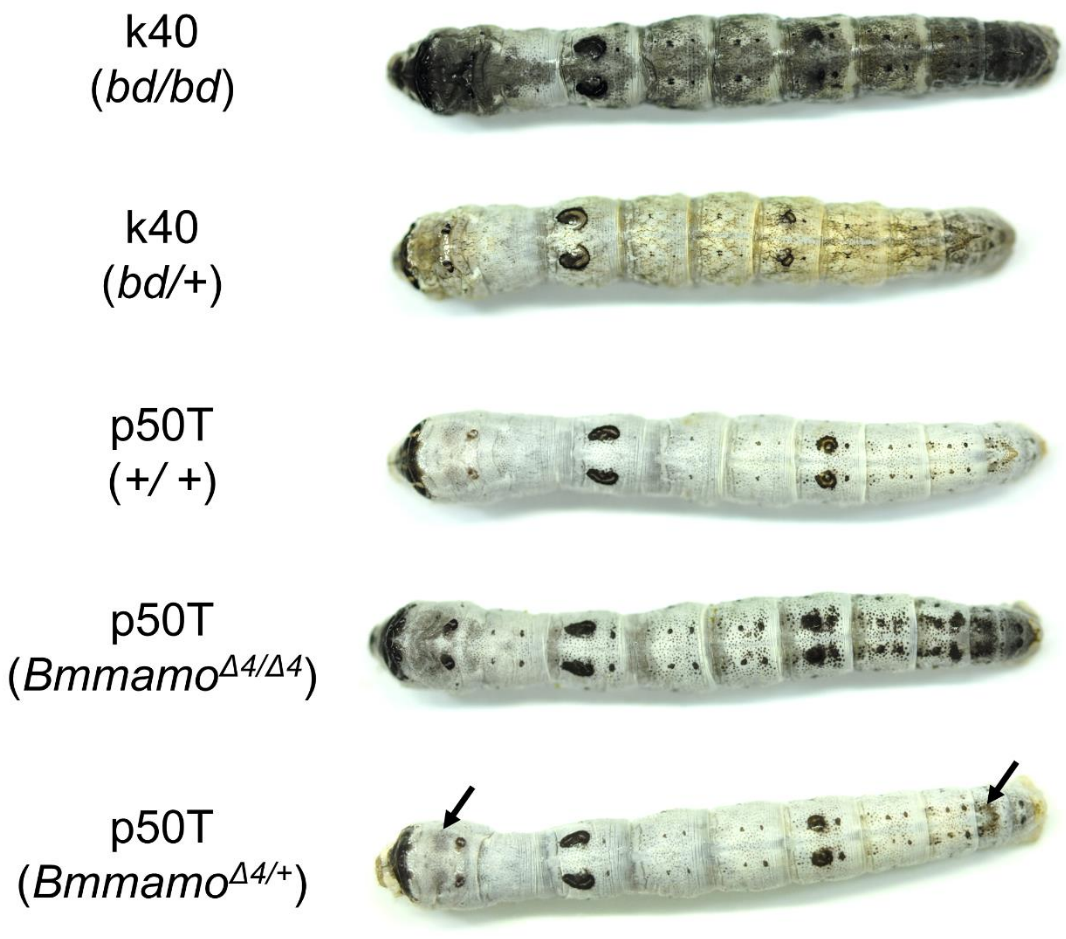


**Figure S6**. Larval phenotypes of k40 strain (*bd/bd* and *bd/*+), p50T strain (+/+), and *Bmmamo* knockout mutants (*Bmmamo*$^{\Delta4/\Delta4}$ and *Bmmamo*$^{\Delta4/+}$). *Bmmamo* knockout mutants were established from the p50T strain (see Materials and Methods). The *Bmmamo*$^{\Delta4/+}$ larvae exhibited a faint black pigmentation on its integument (black arrows), which was not observed in the wild-type p50T larvae.
